# Proteolytic control of FixT by the Lon protease impacts FixLJ signaling in *Caulobacter crescentus*

**DOI:** 10.1101/2024.02.05.579008

**Authors:** Kubra Yigit, Peter Chien

## Abstract

Responding to changes in oxygen levels is critical for aerobic microbes. In *Caulobacter crescentus*, low oxygen is sensed by the FixL-FixJ two-component system which induces multiple genes, including heme biosynthesis, to accommodate microaerobic conditions. The FixLJ inhibitor FixT is also induced under low oxygen conditions and is degraded by the Lon protease, which together provides negative feedback proposed to adjust FixLJ signaling thresholds during changing conditions. Here, we address if the degradation of FixT by the Lon protease contributes to phenotypic defects associated with loss of Lon. We find that Δ*lon* strains are deficient in FixLJ-dependent heme biosynthesis, consistent with elevated FixT levels as deletion of *fixT* suppresses this defect. Transcriptomics validate this result as there is diminished expression of many FixLJ-activated genes in Δ*lon*. However, no physiological changes in response to microaerobic conditions occurred upon loss of Lon, suggesting that FixT dynamics are not a major contributor to fitness in oxygen limiting conditions. Similarly, stabilization of FixT in Δ*lon* strains does not contribute to any known Lon-related fitness defect, such as cell morphology defects or stress sensitivity. In fact, cells lacking both FixT and Lon are compromised in viability during adaptation to long term aerobic growth. Our work highlights the complexity of protease-dependent regulation of transcription factors and explains the molecular basis of defective heme accumulation in Lon-deficient *Caulobacter*.

**Importance:** The Lon protease shapes protein quality control, signaling pathways, and stress responses in many bacteria species. Loss of Lon often results in multiple phenotypic consequences. In this work, we found a connection between the Lon protease and deficiencies in heme accumulation that then led to our finding of a global change in gene expression due to degradation of a regulator of the hypoxic response. However, loss of degradation of this regulator did not explain other phenotypes associated with Lon deficiencies demonstrating the complex and multiple pathways that this highly conserved protease can impact.

## INTRODUCTION

All cells must adapt to changes in environmental conditions, with much of this adaptation being attributed to changes in gene expression. Regulated degradation by energy-dependent proteases facilitates this adaptation (1) by ensuring the timely degradation of unwanted or misfolded proteins while also actively modulating the abundance of specific regulatory proteins. Through their precise control of protein half-lives, energy-dependent proteases sculpt the landscape of gene expression dynamics (2), influencing cellular responses, adaptation (3), and even the fate of the cell itself (4–8). The Lon protease is a highly conserved ATP-dependent protease that plays a crucial role in protein quality control in both prokaryotic and eukaryotic cells (9, 10). Lon is involved in the response to various stress conditions, such as heat shock (11, 12) and oxidative stress (13, 14) and thought to degrade misfolded or damaged proteins that accumulate under these conditions. The function of Lon in *Caulobacter crescentus* (denoted *Caulobacter* throughout) is multifaceted as it plays a critical role in several processes, including cell cycle regulation⍰⍰ and protein quality control.

The FixLJ pathway in *Caulobacter* regulates gene expression in response to changes in oxygen levels (17), mirroring the regulatory mechanism observed in the control of nitrogen fixation in rhizobia (18, 19). FixL is a membrane-bound histidine kinase that senses changes in oxygen levels (20–22), while FixJ is a response regulator that activates transcription of target genes in response to phosphorylation by FixL (23). When oxygen levels are low, FixL is activated and phosphorylates FixJ (24–26), which then directly activates the transcription of genes involved in adaptation to microaerobic conditions, including heme biosynthesis (17) and indirectly controls expression through controlling transcription factors such as FixK (26). FixK plays a crucial role in activating the expression of genes responsible for the biosynthesis and assembly of terminal oxidases to facilitate efficient oxygen utilization in conditions of low oxygen availability (17, 27, 28) (Fig 1).

**Figure 1.**
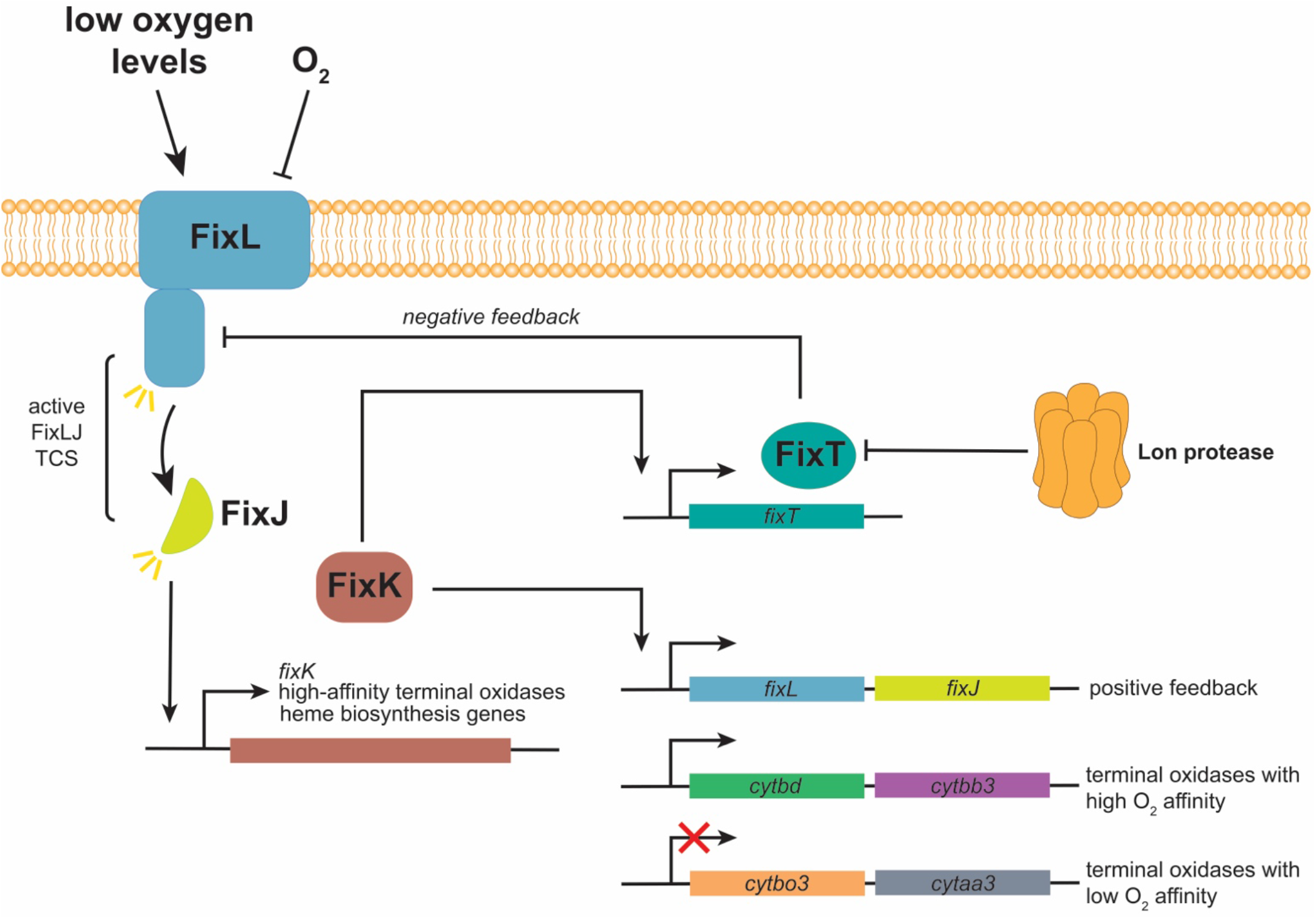
Cartoon depiction of the FixL∼J two-component regulatory system in *Caulobacter crescentus*.

The FixLJ two-component response system in *Caulobacter* is comprised of two feedback regulators: FixK and FixT (17, 29, 30). FixK functions as the positive feedback regulator by turning on the expression of the genes encoding FixL and FixJ. FixK also activates the transcription of the *fixT* gene by binding to its promoter region (17, 31), resulting in the production of the FixT protein. The accumulation of FixT protein leads to inhibition of the autophosphorylation of FixL (29), establishing a negative feedback loop that finely tunes the threshold activity of FixL-FixJ. Interestingly, in *Caulobacter*, FixT is degraded by the Lon protease in the presence of excess oxygen (29). Based on this model, it follows that in Lon-deficient cells the accumulation of the FixT protein should lead to inhibition of the FixLJ pathway and possibly explain some of the pleiotropic phenotypes of a *lon* mutant. Here, we show that the degradation of FixT by the Lon protease controls FixLJ dependent accumulation of heme and aerobic expression of *fixK*, along with other genes in the FixLJ regulon. However, despite this misregulation, stabilization of FixT does not appear to underlie any known damage-sensitivity or physiological defects of Δ*lon* mutants (Fig 1).

## RESULTS

### Deletion of *fixT* restores the accumulation of heme in *lon*-deficient cells

During initial characterization of Δ*lon* strains of *Caulobacter* we noticed that wild type colonies accumulate a red pigment upon prolonged growth on PYE agar plates supplemented with xylose, but Δ*lon* strains do not (Fig 2A). Absorbance spectra of lysates from wild type cultures upon extended growth show a prominent peak at 418 nm (Fig 2B), which corresponds to a Soret maximum, a characteristic signature of molecules containing porphyrin rings and for heme-containing proteins (32). These results suggested that the loss of red pigment in Δ*lon* is reflecting a decrease in heme accumulation. In *Caulobacter*, the FixLJ pathway controls the production of heme by regulating the transcription of the first and sixth enzymes in the heme biosynthetic pathway (33), *hemA* and *hemN*, respectively.

**Figure 2.**
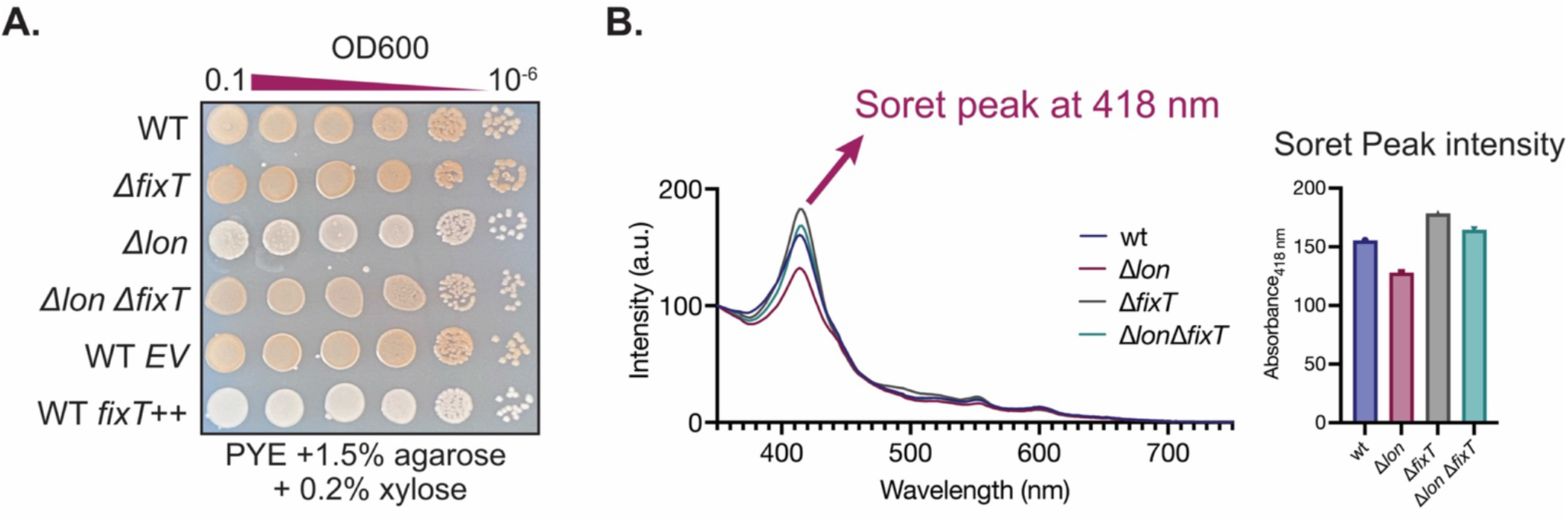
Loss of *fixT* restores the heme accumulation in *lon*-deficient cells. **A**. Serial dilution assays to compare the red pigmentation of WT, Δ*fixT*, Δ*lon*, Δ*fixT*, WT + empty vector (EV) and WT + *fixT*++ strains in PYE agarose plate supplemented with xylose. Cells were spotted to PYE agar plates in 10-fold serial dilutions from left to right. **B**. Left, absorption spectra of the lysates from WT, Δ*lon*, Δ*fixT* and Δ*lon* Δ*fixT* cells with Soret peaks at 418 nm. Right, quantification of the absorption of lysates at 418 nm.

Because recent work showed that a negative regulator of this pathway, FixT, is degraded by the Lon protease in an oxygen dependent manner (29), we reasoned that Δ*lon* strains show reduced accumulation of heme likely due to excess FixT. This would result in repression of FixLJ pathway and the subsequent reduction in heme biosynthesis. To test our hypothesis, we knocked out *fixT* gene in the Δ*lon* strain and found that red pigmentation on plates was restored (Fig 2A) as was heme accumulation based on spectral analysis of the lysates (Fig 2B). Consistent with the role of FixT controlling heme synthesis, overexpressing the *fixT* gene leads to a loss of the red pigmentation phenotype in wild type cells (Fig 2A). Our findings support the model that Lon plays an important role in regulating the FixLJ network through degrading FixT.

### *fixK* expression in Δ*lon* cells is lower than in wild type cells

The FixLJ network becomes active in microaerobic conditions, leading to increased transcription of *fixK*, which subsequently triggers the expression of *fixT* (17, 30). As FixT acts as an inhibitor of FixL autophosphorylation (29), creating a negative feedback loop, we proposed that the elevated levels of FixT protein in Lon-deficient cells should diminish the FixLJ-dependent expression of genes. We analyzed RNA-seq data (34) to identify genes that showed significant changes in the Δ*lon* strain compared to the wild type and compared with previously published microarray data (17) to explore whether genes upregulated by FixL exhibited downregulation in Δ*lon* strains consistent with a surplus of FixT protein (Fig 3A, Table S2).

**Figure 3.**
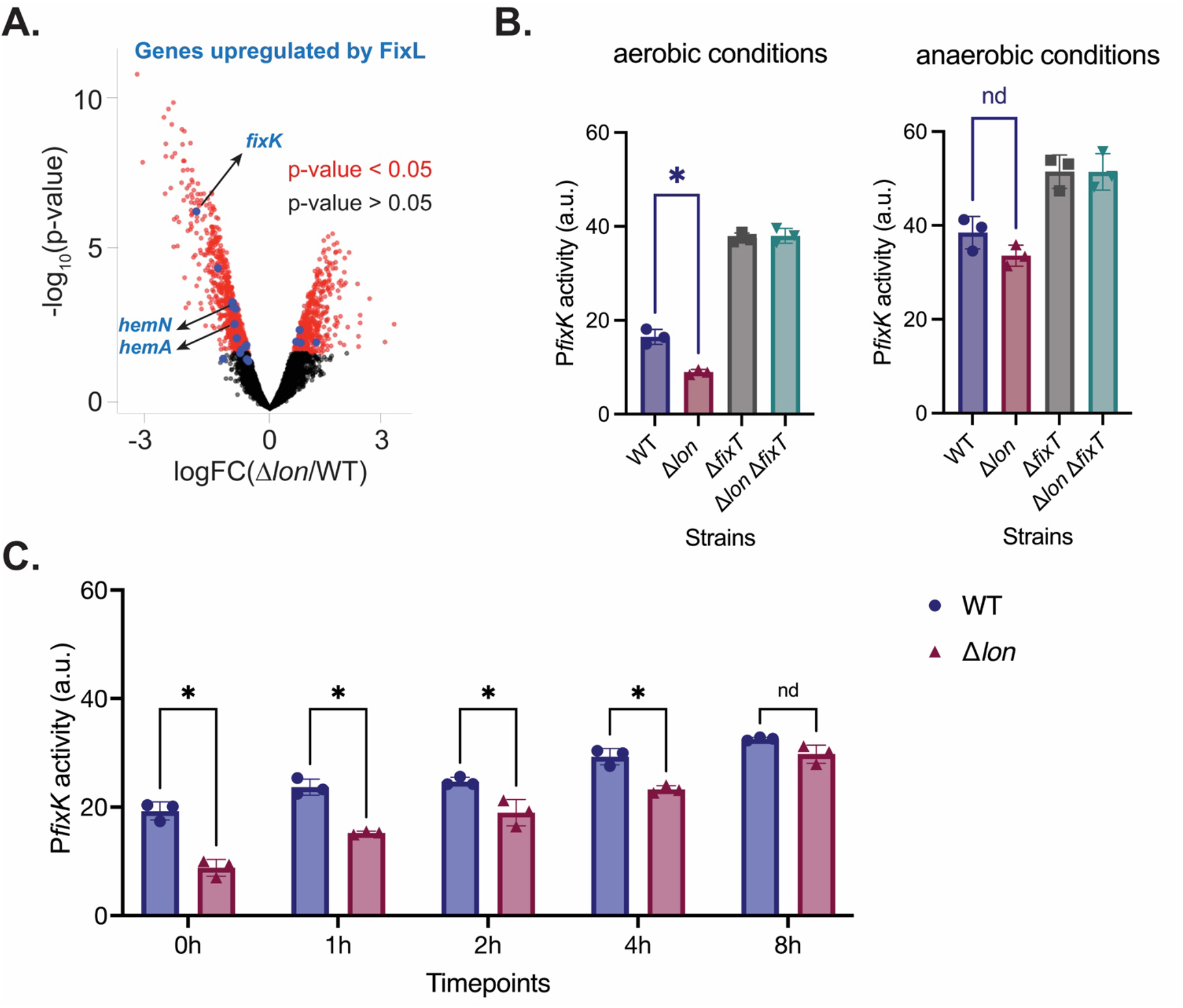
FixT overabundance in Δ*lon* cells represses the expression of *fixK* during aerobic growth and but not in response to hypoxia. **A**. RNA-seq results of WT versus Δ*lon* cells. Red dots represent the genes that are differentially regulated in Δ*lon* compared to WT. Blue dots are the genes that are downregulated upon loss of *fixL*. **B**. pRKlac290-P*fixK* transcriptional reporter activity in WT, Δ*lon*, Δ*fixT* and Δ*lon* Δ*fixT* cells grown under aerobic and microaerobic growth conditions (18hrs growth, average ± SD, n = 3 biological replicates). ^*^, p < 0.05, nd, p > 0.05, t test. **C**. P*fixK*-lacZ reporter activity at different timepoints in WT and Δ*lon* cells grown in PYE with no aeration (average ± SD, n = 3 biological replicates). ^*^, p < 0.05, nd, p > 0.05, t test.

Our examination revealed a clear trend: genes that are typically activated by FixL exhibited significantly diminished expression levels in Δ*lon* strains (Fig 3A, Table S2). Particularly noteworthy were the downregulation of *hemA* and *hemN*, which explains the heme accumulation phenotype, along with a significant reduction in the expression of *fixK*. To validate the effects on *fixK* expression, we employed a beta-galactosidase based transcriptional reporter, where *lacZ* expression is controlled by the *fixK* promoter (P*fixK)*. Based on this assay, we found a two-fold decrease in activity in Δ*lon* cells compared to wild type cells under aerobic conditions (Fig 3B). Following *fixT* deletion, the *fixK* reporter activity increased two-fold in wild type strains and five-fold in Δ*lon* strains, eliminating the differences in reporter expression between WT and Δ*lon* strains (Fig 3B). These results strongly suggest that Lon regulates *fixK* expression through a mechanism dependent on FixT. Because FixLJ is activated during oxygen limitation, we were pleased to see the expected increase in *fixK* promoter activity during microaerobic growth conditions in both WT and Δ*lon* (Fig 3C). Interestingly, when we shift aerobically grown cells to microaerobic conditions, *fixK* promoter activity was induced proportionally more in the Δ*lon* (300% increase) than the WT (50% increase), with negligible differences in promoter activity once cells fully adapted to microaerobic conditions (Fig 3C). We conclude that excess FixT suppression of FixLJ signaling is most impactful during oxygen replete conditions and that other regulatory pathways supersede this suppression when cells fully adapt to oxygen limiting conditions.

### FixT affects growth under changing oxygen conditions only in the absence of Lon

We next explored whether destabilization of FixT by Lon resulted in defects during growth under oxygen changing conditions. To test this, we grew cells in either oxygen limiting or standard aerobic conditions, then measured growth kinetics when cells were shifted to standard aerobic conditions. Following standard aerobic growth overnight, Δ*lon* cells accumulate less cell mass upon entry to stationary phase (34), an effect that was not rescued by deletion of *fixT* (Fig 4A). Interestingly, while the loss of *fixT* was well tolerated in wild type cells, deletion of *fixT* led to a slower growth in Δ*lon* strains (Fig 4A).

**Figure 4.**
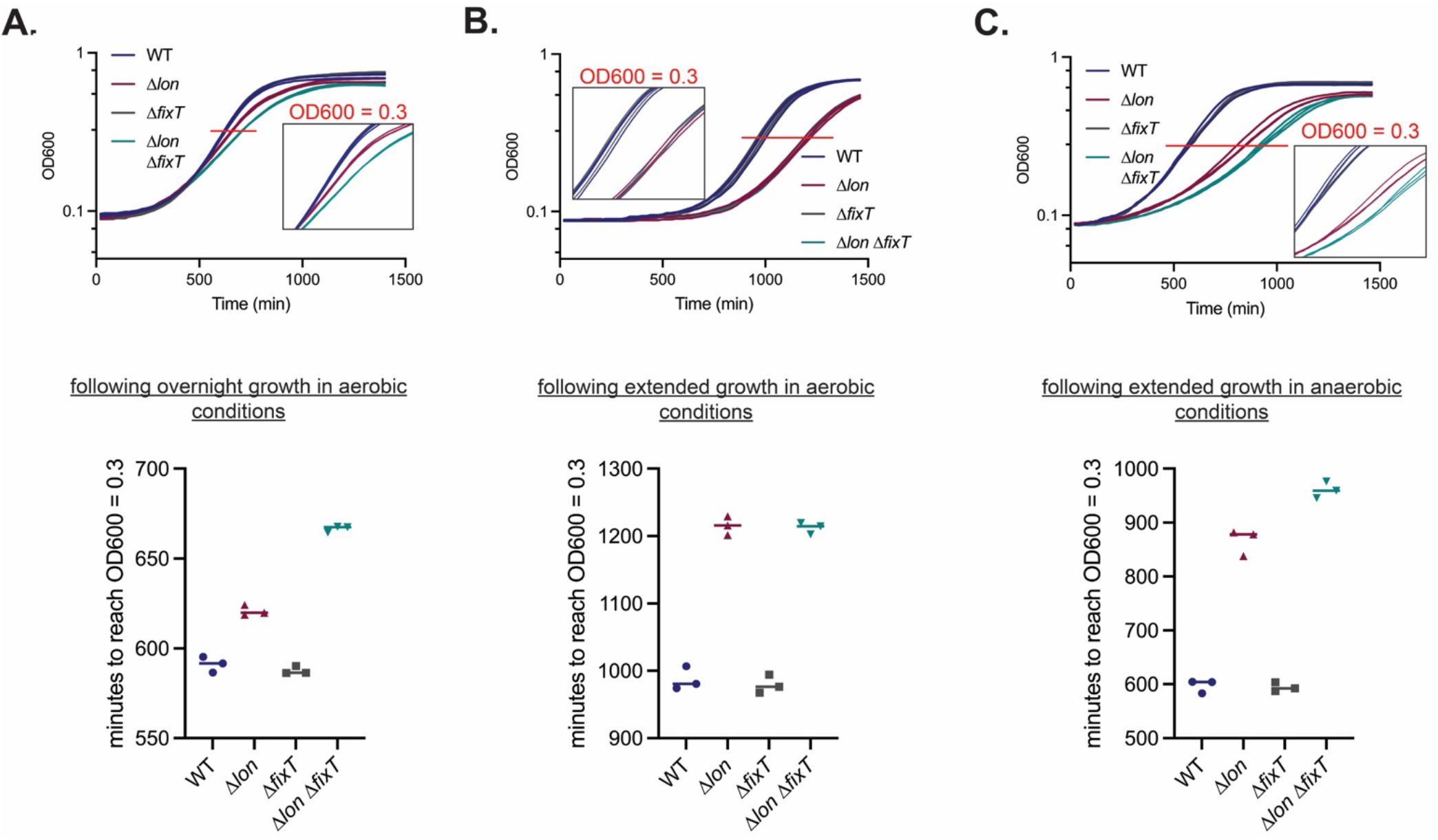
**A.** Top, 24h growth curve of WT, Δ*fixT*, Δ*lon* and Δ*lon* Δ*fixT* cells following an overnight incubation in PYE media. Bottom, graph showing the time it takes for WT, Δ*fixT*, Δ*lon* and Δ*lon* Δ*fixT* cells grown overnight in PYE media to reach an OD600 of 0.3 (n = 3 biological replicates). **B**. Top, 24h growth curve of WT, Δ*fixT*, Δ*lon* and Δ*lon* Δ*fixT* cells in fresh PYE media after 10 days of growth in PYE media under aerobic conditions. Bottom, graph showing the time it takes for WT, Δ*fixT*, Δ*lon* and Δ*lon* Δ*fixT* cells to reach an OD600 of 0.3 during recovery from prolonged aerobic growth in PYE (n = 3 biological replicates). **C**. 24h growth curve of WT, Δ*fixT*, Δ*lon* and Δ*lon* Δ*fixT* cells in fresh PYE media after 10 days of growth in PYE media under microaerobic conditions. Bottom, graph showing the time it takes for WT, Δ*fixT*, Δ*lon* and Δ*lon* Δ*fixT* cells to reach an OD600 of 0.3 during recovery from prolonged microaerobic growth in PYE (n = 3 biological replicates).

Compared to standard aerobic growth, regrowth of wild type cells shifted from prolonged aerobic conditions to standard aerobic conditions resulted in cells taking longer to exit the lag phase, an effect even more pronounced in Δ*lon* (Fig 4B). By contrast, wild type cells exiting prolonged microaerobic conditions begin to grow rapidly when reoxygenated while Δ*lon* strains have defects with this regrowth (Fig 4C). Interestingly, when both *lon* and *fixT* are deleted, this growth defect is further accentuated (Fig 4C). Overall, the loss of FixT consistently led to a growth defect in Δ*lon* cells, regardless of oxygen levels or growth duration.

### Excess FixT does not explain other lon-associated phenotypes

The absence of the Lon protease is associated with various physiological abnormalities during standard growth conditions, including reduced growth rate, diminished growth on low-percentage agar, cell filamentation, and elongated stalk lengths (34–36). Given that reduced heme pools in *lon* deletions are caused by elevated FixT levels, we investigated whether FixT stabilization could explain other Δ*lon*-related phenotypes. Microscopic analysis of wild type, Δ*lon*, Δ*fixT*, and Δ*lon* Δ*fixT* cells in the exponential growth phase show that absence of *fixT* does not influence cell morphology nor does it rescue the filamentation phenotype of Δ*lon* (Fig 5A). Similarly, the reduced growth observed in Δ*lon* mutants on low-percentage agar, which reflects a combination of motility and growth, remains the same in Δ*lon* Δ*fixT* strain (Fig 5B). *Caulobacter lon* deletion mutants show heightened sensitivity to various stressors, including DNA damage induced by agents like mitomycin C (MMC) (35) and oxidative stress caused by hydrogen peroxide (H_2_O_2_) *and* Δ*lon* Δ*fixT* are as sensitive to these stresses as Δ*lon* (Fig 5C, D). We conclude that a surplus of FixT is not solely responsible for any of the stress-related or morphologically aberrant features of Δ*lon* strains.

**Figure 5.**
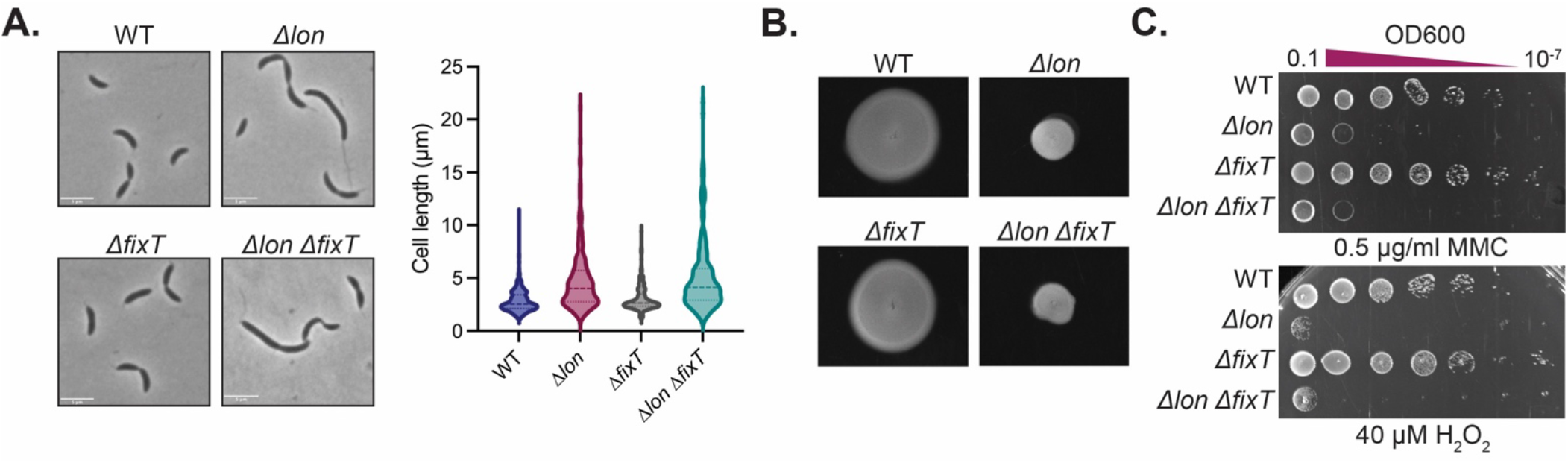
*fixT* deletion does not rescue physiology defects of Lon-deficient *Caulobacter* cells under laboratory conditions. **A**. Left, representative phase-contrast microscopy images of WT, ΔfixT, Δ*lon* and Δ*lon* Δ*fixT* cells grown in PYE visualized at exponential phase. Scale bar = 5 µm. Right, quantification of the length of cells from each strain. **B**. WT, Δ*fixT*, Δ*lon* and Δ*lon* Δ*fixT* single colonies grown in motility agar (0.3% PYE agar). **C**. Serial dilution assays to compare the viability of strains in PYE agar plates supplemented with either 0.5 µg/mL mitomycin C (MMC) to induce genotoxic stress or 40 µM hydrogen peroxide (H_2_O_2_) to generate oxidative stress. Cells were spotted to PYE agar plates in 10-fold serial dilutions from left to right.

## DISCUSSION

In *Caulobacter* cells lacking the Lon protease (Δ*lon*) there is an intracellular surplus of the FixT protein due to lack of degradation (29). This excess FixT exerts inhibitory effects on the FixLJ two-component system and its downstream regulon, including the heme biosynthetic pathway. Our studies revealed that the red pigmentation of *Caulobacter* colonies on xylose-containing PYE plates is the result of heme accumulation, which is influenced by control of FixT by the Lon protease (Fig 2A, B). In addition, our comparative analysis of the RNA-Seq and previously published microarray datasets showed that the absence of Lon results in the downregulation of genes normally activated by FixL, which we confirm with reporter assays and includes heme biosynthesis genes. We conclude that Lon plays an important role in managing the FixT-dependent feedback control of the FixLJ regulon. However, the disappearance of this difference in oxygen-limiting conditions indicate that the elevated levels of FixT in Δ*lon* cells no longer suppress the FixLJ regulon in those conditions (Fig 6).

**Figure 6.**
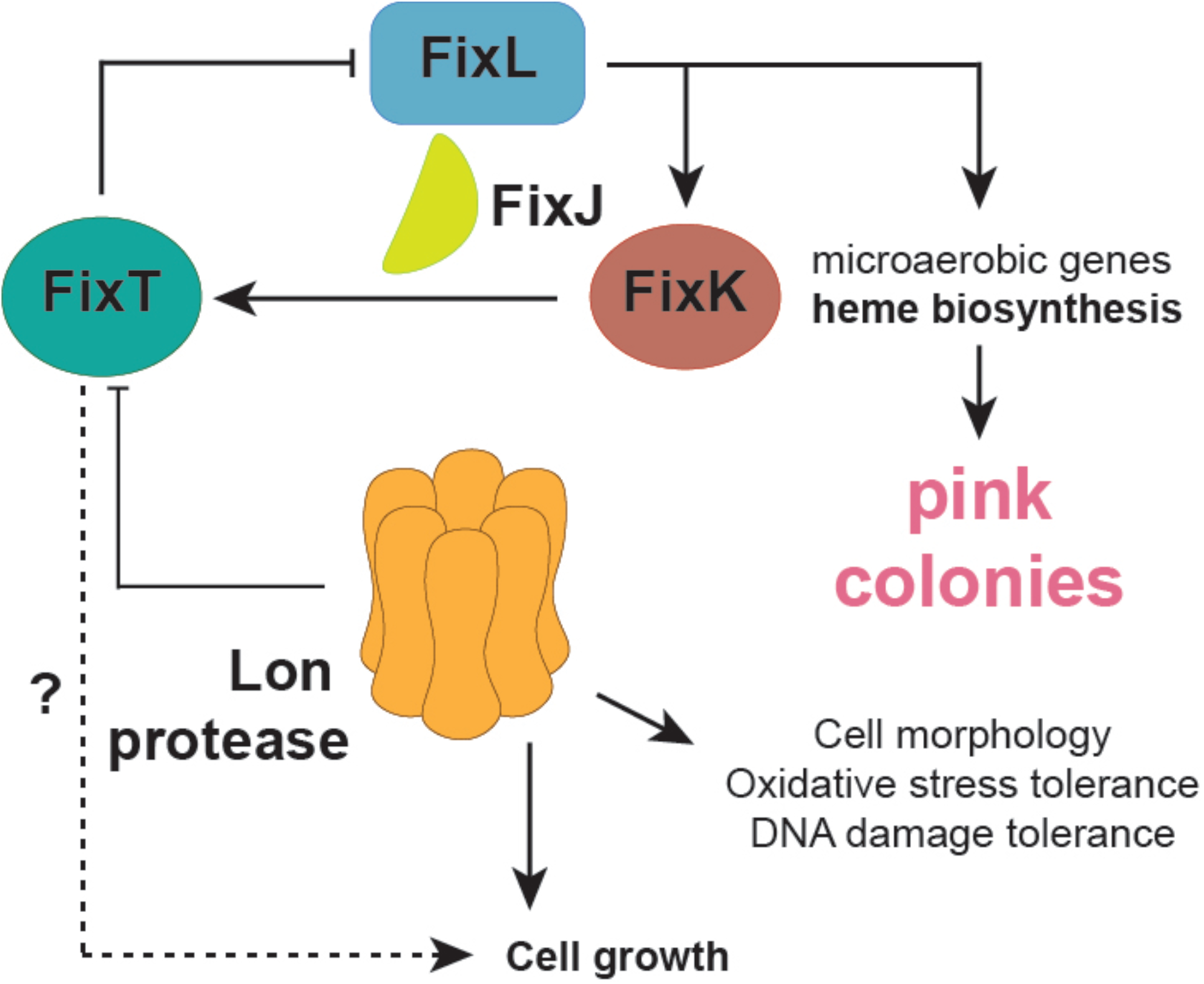
Our model of how the regulation of FixT through Lon-mediated proteolysis affects *Caulobacter* physiology. Lon’s degradation of FixT in aerobic growth conditions results in an overabundance of FixT in the absence of Lon. This overabundance most likely results in the suppression of FixL activity and subsequently, as our results show, a reduction of *fixK* expression and the heme production. FixT accumulation in Δ*lon* cells however does not contribute to any of the *lon* phenotypes including cell motility and morphology, sensitivity to DNA damaging agents or oxidative stress. Interestingly, deletion of *fixT* in Δ*lon* cells leads to slower growth, which indicates a synergistic effect between Lon and FixT in controlling cell growth.

Regrowth experiments under different conditions reveal that the loss of *fixT* results in a prolonged growth delay in Δ*lon* cells but not in wild type cells (Fig 4A) suggesting that Lon and FixT play independent roles in controlling *Caulobacter* cell growth and the deletion of both causes a synergistic effect. These effects are oxygen and cell phase dependent as prolonged growth in aerobic conditions suppresses the deleterious consequences seen when both FixT and Lon are absent in prolonged microaerobic or single overnight growth conditions (Fig 4B, C). We speculate that either imbalances in general protein homeostasis in the cell due to loss of Lon or additional factors regulated by Lon impinge on similar downstream targets as those FixT regulates (likely through the FixLJ controlled pathway) as depicted in Figure 6.

Finally, our investigation into other Lon-related phenotypes did not reveal any evidence for FixT degradation by Lon playing a significant role in *Caulobacter* physiology in laboratory conditions (Fig 5A, B, C). Supporting this statement are our observations that despite the visible defects in heme accumulations, Δ*lon* Δ*fixT* strains are phenotypically identical to Δlon strains for all known Lon-related defects. Thus, if FixT proteolysis by Lon has some physiological consequence, it relies on growth conditions or environmental stresses that are currently unknown.

## MATERIALS AND METHODS

### Bacterial strains and growth conditions

Please refer to Table S1 for detailed information on plasmids and strains. All the *Caulobacter crescentus* strains utilized in this research originated from the NA1000 strain. We gratefully acknowledge the Crosson Lab for providing the plasmids used in this study. Liquid cultures of *Caulobacter crescentus* strains were cultivated in peptone yeast extract (PYE) medium, which comprised 2 g/L peptone, 1 g/L yeast extract, 1 mM MgSO_4_, and 0.5 mM CaCl_2_. For solid cultures, the cells were grown on PYE media containing 1.5% agar. The *lon* clean deletion strain was previously established (34). *fixT* knockout strain was generated in wild type NA1000 strain through a two-step recombination method using the pNPTS138-Δ*fixT* plasmid. Wild type *Caulobacter* cells were transformed with the pNPTS138-Δ*fixT* plasmid via electroporation and the initial selection was conducted on PYE agar plates containing kanamycin (0.25 mg/mL). Colonies displaying successful primary integrations were chosen and cultured overnight in PYE media. Subsequently, they were plated on PYE agar plates containing 2% w/v sucrose to assess whether the plasmid sequence had been eliminated from the chromosome through secondary recombination. After the sucrose selection, colonies were patched on PYE agar plates, with or without kanamycin. The loss of kanamycin resistance served as an indicator of plasmid removal. To confirm the deletion, colony PCR was carried out on the cells that did not grow on kanamycin, using forward and reverse primers targeting the *fixT* coding sequence. Colonies lacking the *fixT* sequence were further verified through whole genome sequencing. To generate the Δ*lon* Δ*fixT* strain, *lon:gentR* was transduced to Δ*fixT* NA1000 cells using the phiCR30 phage. For overexpressing *fixT in vivo*, we introduced the pMT805-*fixT* plasmid into the NA1000 strain via electroporation, with subsequent selection on PYE agar plates containing chloramphenicol (0.02 mg/mL). An empty pMT805 plasmid was employed as a control. Liquid cultures were then cultivated in PYE media supplemented with chloramphenicol (0.01 mg/mL).

### Whole genome sequencing

Genomic DNA was isolated from the cells using Monarch^®^ Genomic DNA Purification Kit (New England Biolabs). DNA concentration was measured with Nanodrop One/One^C^ spectrophotometer (Thermo Fisher). Genomic DNA extracted from the cells were sent to SeqCenter, LLC (Pittsburgh, PA) for Illumina whole genome sequencing. The FASTQ files were analyzed using breseq (37) to confirm the gene knockouts.

### Phenotypic analysis

Strains were grown overnight in 10 ml PYE to saturation at 30 ºC and back diluted to OD600 of 0.1 and grown to mid exponential phase. Prior to plating, the cells were normalized to OD600 of 0.1 and 10-fold serially diluted in PYE media in a 96-well plate. For the analysis of red pigmentation, PYE agarose plates containing 0.2% xylose were prepared. 3 µL of serially diluted cells were transferred from the wells of the 96-well plate by a multichannel pipette to the xylose-containing PYE-agarose plate. For the spectral analyses of heme content, equal number of cells from stationary phase cultures were centrifuged and lysed in BugBuster lysis reagent diluted to 1X in H-Buffer (20 mM HEPES, 100 mM KCl, 10 mM MgCl_2_.6H_2_O, 10% glycerol, pH 7.5). Concentration of the proteins in the lysates were measured by Pierce™ Coomassie (Bradford) Protein Assay Reagent. Protein concentration in the lysates were normalized to 10 mg/ml and the lysates were transferred to 96-well clear bottom culture plates. The absorbance was read from 350 nm to 750 nm in SpectraMax M2 plate reader (Molecular Devices).

### RNA-seq and ß-galactosidase assays

For gene expression analyses, previously published RNA-seq results (34) and microarray datasets (17) were used. In Table S1, enrichment of FixL, FixJ and FixK regulated genes in Δlon differentially regulated expression was determined with fry analysis (edgeR).

For beta-galactosidase assays, *Caulobacter* cells were transformed with pRKlac290-P*ctrA* and pRKlac290-P*fixK* plasmids via electroporation and selected on PYE agar plates containing oxytetracycline (2 µg/mL). Liquid cultures were grown in PYE media supplemented with oxytetracycline (1 µg/mL. 15 mg/mL tetracycline stock solution was prepared in 70% ethanol. To assess *fixK* and *ctrA* (control) promoter activity under different conditions, overnight cultures were back diluted to OD600 = 0.05 and allowed to grow until reaching an OD600 of 0.4. For the analysis of aerobic *fixK* expression, cells were lysed immediately and for the microaerobic expression analysis, 5 mL from each culture was transferred to 5 mL culture tubes and the tubes were sealed tightly. The tubes were positioned upright in a 30 ºC shaker to minimize the oxygenation of the growth media. Cells were lysed after ∼18 hours of microaerobic growth. For timepoint assays, overnight cultures were back diluted to OD600 = 0.05 and grown until OD600 = 0.4. Cells that were lysed immediately were considered the initial hour samples and the rest of the cells were then transferred to 5 mL culture tubes and grown in microaerobic-mimicking conditions. Four tubes of the same culture were prepared for four timepoints as the hourly sample collection prevents the depletion of oxygen in the tubes. Cells were collected and lysed at each time point and tested for beta-galactosidase activity. Cell lysis was achieved via chloroform-permeabilization in 96-well 2.2 mL polypropylene blocks containing 1 mL Z-buffer (60 mM Na_2_HPO_4_, 40 mM NaH_2_PO_4_, 10 mM KCl, 1 mM MgSO_4_), 20 µL 0.1% w/v SDS and 40 µL chloroform (38). Following the addition of cells to the wells, the cells were lysed by pipetting the liquid inside the wells for 15-20 times. After the chloroform in the wells reached the bottom, 100 µL lysate was collected from the surface of each well and transferred to a clear 96-well flat bottom plate. 20 µL of ONPG solution (4 mg/mL in distilled water) was added to each well and the absorbance at 420 nm was measured for 40 minutes in SpectraMax M2 plate reader (Molecular Devices). The reaction was quenched with the addition of 50 µL 1M Na_2_CO_3_.

### Bacterial growth curves

Cells were grown overnight in 30 °C shaking incubator and back diluted to OD600 = 0.025. Upon reaching an OD600 of 0.4, the cells were transferred to a clear, flat bottom 96-well plate and diluted to a starting OD600 of 0.0025 in PYE to assess the regrowth kinetics after an overnight growth. To mimic microaerobic conditions, 14 mL cultures at OD600 = 0.4 were transferred to 14-mL culture tubes and the tubes were sealed by pushing the cap tightly. Tubes were placed in 30 ºC shaker in a standing-up position, not tilted, and cells were allowed to grow for 4 days. The prolonged aerobic growth of cells was achieved by transferring 1.4 mL of exponential phase cultures to 14-mL culture tubes without sealing and the cells were grown for 4 days in a 30 °C shaking incubator. Regrowth kinetics after prolonged microaerobic and aerobic growth was assessed by transferring the cells to a clear flat bottom 96-well plate with a starting OD600 of 0.0025. The 96-well plates were placed in Biotek Epoch 2 Microplate Spectrophotometer (Agilent). OD600 measurements were made for 24 hours in 20-minute intervals and the OD600 values were plotted in GraphPad Prism software.

### Microscopy, cell morphology and viability assays

Replicates of cells were grown overnight in PYE media and back diluted to OD600 = 0.025. 3 µL mid-exponential phase cultures were mounted on 1.5% PYE agarose pads. Phase contrast images of the exponential phase cultures were taken using Zeiss AXIO Scope A1. Cell length was measured using MicrobeJ plugin of the ImageJ software (Fiji). 200 cells were counted for each strain and the lengths were plotted in GraphPad Prism. Motility analyses were done by placing a single colony (n = 3) of each strain to soft agar plates (PYE + 0.3% agar). Plates were places in a 30 °C incubator and colonies were allowed to spread for 2 days. Plates were imaged with SynGene G:Box iChemi XT Imager. For viability assays, 400 µg/mL mitomycin C (Millipore Sigma) stock solution was prepared in distilled water. 3% (w/v) hydrogen peroxide (H_2_O_2_) was freshly diluted to 10 mM before the experiment. To determine MMC and H_2_O_2_ sensitivity, PYE agar plates supplemented with 0.5 µg/mL mitomycin C and 40 µM H_2_O_2_ were prepared. Overnight cultures were back diluted to OD600 = 0.025 and allowed to grow to mid-exponential phase prior to 10-fold serial dilution and plating on mitomycin C containing PYE agar plates. For H_2_O_2_ sensitivity, overnight cultures were serially diluted and spotted on H_2_O_2_ containing PYE agar plates. Plates were transferred to a 30 °C incubator for growth and pictures were taken after 2 days using Syngene G:Box.

## Acknowledgments

This project was supported by NIH R35GM130320 to Peter Chien. K. Y. was supported in part through the University of Massachusetts Amherst NIH Chemistry-Biology Interface Training Program (NIH T32GM139789). The pNPTS138-Δ*fixT*, pMT805-*fixT* and pRKlac290-PfixK plasmids were generously gifted by Sean Crosson (Michigan State University). We thank Rilee D. Zeinert and Kethney Massenat for the initial characterization of the red pigmentation, and Chien lab members and Sean Crosson for helpful discussion.

---

**Supplemental Table 1.** Plasmids and *Caulobacter* strains used in this study.

| Plasmid or strain | Relevant characteristics | Comments | Reference |
| --- | --- | --- | --- |
| NA1000 | Wild type strain |  | This study |
| CPC667 | NA1000 / $\Delta lon$ | | (39) |
| CPC1225 | NA1000 / $\Delta fixT$ | | This study |
| CPC1226 | NA1000 / $\Delta lon$<br>$\Delta fixT$ | | This study |
| FC106 | pNPTS138-$\Delta fixT$ | Allele replacement plasmid for deletion of *fixT* | (34) |
| EPC407 | pMT805 | Replicating plasmid (mid copy, BBR origin) with xylose inducible promoter, Chlor<sup>R</sup> | This study |
| FC3396 | pMT805-*fixT* | Replicating plasmid containing *fixT* coding sequence (mid copy, BBR origin) with xylose inducible promoter, Chlor<sup>R</sup> | (34) |
| EPC615 | pRKlac290-*PctrA* | Plasmid containing *PctrA*-lacZ transcriptional fusion, Tet<sup>R</sup> | This study |
| FC94 | pRKlac290-*PfixK* | Plasmid containing *PfixK*-lacZ transcriptional fusion, Tet<sup>R</sup> | (34) |

**Supplemental Table 2. Analysis of microarray and RNA sequencing results**.

## References

1. Mahmoud SA, Chien P. 2018. Regulated Proteolysis in Bacteria. Annu Rev Biochem 87:1– 20.

2. Dyck L van, Neupert W, Langer T. 1998. The ATP-dependent PIM1 protease is required for the expression of intron-containing genes in mitochondria. Genes Dev 12:1515–1524.

3. Wettstadt S, Llamas MA. 2020. Role of Regulated Proteolysis in the Communication of Bacteria With the Environment. Front Mol Biosci 7:586497.

4. Konovalova A, Søgaard-Andersen L, Kroos L. 2014. Regulated proteolysis in bacterial development. FEMS Microbiol Rev 38:493–522.

5. Ehrmann M, Clausen T. 2004. PROTEOLYSIS AS A REGULATORY MECHANISM. Genetics 38:709–724.

6. Lau J, Hernandez-Alicea L, Vass RH, Chien P. 2015. A Phosphosignaling Adaptor Primes the AAA+ Protease ClpXP to Drive Cell Cycle-Regulated Proteolysis. Mol Cell 59:104–116.

7. Joshi KK, Bergé M, Radhakrishnan SK, Viollier PH, Chien P. 2015. An Adaptor Hierarchy Regulates Proteolysis during a Bacterial Cell Cycle. Cell 163:419–431.

8. Gora KG, Cantin A, Wohlever M, Joshi KK, Perchuk BS, Chien P, Laub MT. 2013. Regulated proteolysis of a transcription factor complex is critical to cell cycle progression in Caulobacter crescentus. Mol Microbiol 87:1277–89.

9. Lee I, Suzuki CK. 2008. Functional mechanics of the ATP-dependent Lon protease-lessons from endogenous protein and synthetic peptide substrates. Biochim Biophys Acta (BBA) - Proteins Proteom 1784:727–735.

10. Gur E, Sauer RT. 2008. Recognition of misfolded proteins by Lon, a AAA+ protease. Genes Dev 22:2267–2277.

11. Phillips TA, VanBogelen RA, Neidhardt FC. 1984. lon gene product of Escherichia coli is a heat-shock protein. J Bacteriol 159:283–287.

12. Tsilibaris V, Maenhaut-Michel G, Melderen LV. 2006. Biological roles of the Lon ATP-dependent protease. Res Microbiol 157:701–713.

13. Bota DA, Davies KJA. 2002. Lon protease preferentially degrades oxidized mitochondrial aconitase by an ATP-stimulated mechanism. Nat Cell Biol 4:674–680.

14. Ngo JK, Davies KJA. 2009. Mitochondrial Lon protease is a human stress protein. Free Radic Biol Med 46:1042–1048.

15. Jenal U. 2009. The role of proteolysis in the Caulobacter crescentus cell cycle and development. Res Microbiol 160:687–695.

16. Jonas K, Liu J, Chien P, Laub MT. 2013. Proteotoxic Stress Induces a Cell-Cycle Arrest by Stimulating Lon to Degrade the Replication Initiator DnaA. Cell 154:623–636.

17. Crosson S, McGrath PT, Stephens C, McAdams HH, Shapiro L. 2005. Conserved modular design of an oxygen sensory/signaling network with species-specific output. Proc Natl Acad Sci 102:8018–8023.

18. David M, Daveran M-L, Batut J, Dedieu A, Domergue O, Ghai J, Hertig C, Boistard P, Kahn D. 1988. Cascade regulation of nif gene expression in Rhizobium meliloti. Cell 54:671–683.

19. Agron PG, Monson EK, Ditta GS, Helinski DR. 1994. Oxygen regulation of expression of nitrogen fixation genes in Rhizobium meliloti. Res Microbiol 145:454–459.

20. Gilles-Gonzalez MA, Ditta GS, Helinski DR. 1991. A haemoprotein with kinase activity encoded by the oxygen sensor of Rhizobium meliloti. Nature 350:170–172.

21. Philip P de, Batut J, Boistard P. 1990. Rhizobium meliloti Fix L is an oxygen sensor and regulates R. meliloti nifA and fixK genes differently in Escherichia coli. J Bacteriol 172:4255– 4262.

22. Perutz MF, Paoli M, Lesk AM. 1999. Fix L, a haemoglobin that acts as an oxygen sensor: signalling mechanism and structural basis of its homology with PAS domains. Chem Biol 6:R291–R297.

23. Fischer HM. 1994. Genetic regulation of nitrogen fixation in rhizobia. Microbiol Rev 58:352– 386.

24. Tuckerman JR, Gonzalez G, Dioum EM, Gilles-Gonzalez M-A. 2002. Ligand and Oxidation-State Specific Regulation of the Heme-Based Oxygen Sensor FixL from Sinorhizobium meliloti †. Biochemistry 41:6170–6177.

25. Dixon R, Kahn D. 2004. Genetic regulation of biological nitrogen fixation. Nat Rev Microbiol 2:621–631.

26. Bobik C, Meilhoc E, Batut J. 2006. FixJ: a Major Regulator of the Oxygen Limitation Response and Late Symbiotic Functions of Sinorhizobium meliloti. J Bacteriol 188:4890–4902.

27. Bueno E, Mesa S, Bedmar EJ, Richardson DJ, Delgado MJ. 2012. Bacterial Adaptation of Respiration from Oxic to Microoxic and Anoxic Conditions: Redox Control. Antioxid Redox Signal 16:819–852.

28. Gabel C, Maier RJ. 1993. Oxygen-dependent transcriptional regulation of cytochrome aa3 in Bradyrhizobium japonicum. J Bacteriol 175:128–132.

29. Stein BJ, Fiebig A, Crosson S. 2020. Feedback control of a two-component signaling system by an Fe-S-binding receiver domain. mBio 729053.

30. Foussard M, Garnerone A, Ni F, Soupène E, Boistard P, Batut J. 1997. Negative autoregulation of the Rhizobium meliloti fixK gene is indirect and requires a newly identified regulator, FixT. Mol Microbiol 25:27–37.

31. Garnerone A-M, Cabanes D, Foussard M, Boistard P, Batut J. 1999. Inhibition of the FixL Sensor Kinase by the FixT Protein inSinorhizobium meliloti ^*^. J Biol Chem 274:32500–32506.

32. Luthra A, Denisov IG, Sligar SG. 2011. Spectroscopic features of cytochrome P450 reaction intermediates. Arch Biochem Biophys 507:26–35.

33. Dailey HA, Dailey TA, Gerdes S, Jahn D, Jahn M, O’Brian MR, Warren MJ. 2017. Prokaryotic Heme Biosynthesis: Multiple Pathways to a Common Essential Product. Microbiol Mol Biol Rev 81.

34. Mahmoud SA, Aldikacti B, Chien P. 2022. ATP hydrolysis tunes specificity of a AAA+ protease. Cell Rep 40:111405.

35. Zeinert RD, Baniasadi H, Tu BP, Chien P. 2020. The Lon Protease Links Nucleotide Metabolism with Proteotoxic Stress. Mol Cell 79:758–767.e6.

36. Omnus DJ, Fink MJ, Szwedo K, Jonas K. 2021. The Lon protease temporally restricts polar cell differentiation events during the Caulobacter cell cycle. eLife 10:e73875.

37. Deatherage DE, Barrick JE. 2014. Engineering and Analyzing Multicellular Systems, Methods and Protocols. Methods Mol Biol 1151:165–188.

38. Griffith KL, Wolf RE. 2002. Measuring β-Galactosidase Activity in Bacteria: Cell Growth, Permeabilization, and Enzyme Assays in 96-Well Arrays. Biochem Biophys Res Commun 290:397–402.

